# Secretory IgM (sIgM) is an ancient master regulator of microbiota homeostasis and metabolism

**DOI:** 10.1101/2023.02.26.530119

**Authors:** Yang Ding, Alvaro Fernández-Montero, Amir Mani, Elisa Casadei, Yasuhiro Shibasaki, Fumio Takizawa, Ryuichiro Miyazawa, Irene Salinas, J. Oriol Sunyer

## Abstract

The co-evolution between secretory immunoglobulins (sIgs) and microbiota began with the emergence of IgM over half a billion years ago. Yet, IgM function in vertebrates is mostly associated with systemic immunity against pathogens. sIgA and sIgT are the only sIgs known to be required in the control of microbiota homeostasis in warm- and cold-blooded vertebrates respectively. Recent studies have shown that sIgM coats a large proportion of the gut microbiota of humans and teleost fish, thus suggesting an ancient and conserved relationship between sIgM and microbiota early in vertebrate evolution. To test this hypothesis, we temporarily and selectively depleted IgM from rainbow trout, an old bony fish species. IgM depletion resulted in a drastic reduction in microbiota IgM coating levels and losses in gut-associated bacteria. These were accompanied by bacterial translocation, severe gut tissue damage, inflammation and dysbiosis predictive of metabolic shifts. Furthermore, depletion of IgM resulted in body weight loss and lethality in an experimental colitis model. Recovery of sIgM to physiological levels restores tissue barrier integrity, while microbiome homeostasis and their predictive metabolic capabilities are not fully restituted. Our findings uncover a previously unrecognized role of sIgM as an ancient master regulator of microbiota homeostasis and metabolism and challenge the current paradigm that sIgA and sIgT are the key vertebrate sIgs regulating microbiome homeostasis.

**One-Sentence Summary:** IgM, the most ancient and conserved immunoglobulin in jawed vertebrates, is required for successful symbiosis with the gut microbiota.

## Main Text

The emergence of immunoglobulin (Ig) genes in jawed vertebrates over 500 million years ago marked a milestone in the evolution of adaptive immune systems (*1, 2*). IgM is the oldest, most functionally stable and conserved Ig in all jawed vertebrates (*3*). While IgM function is mostly associated with systemic immune responses to pathogens, IgM can also be found in mucosal secretions of several vertebrates as secretory IgM (sIgM) (*4-6*). Besides sIgM, these mucosal sites contain specialized mucosal Igs (sIgA in mammals and sIgT in fish) that are known to coat a large portion of their microbiota and play a key role in the regulation microbiome homeostasis. Recent observations showing that sIgM can also be found coating a portion of the microbiota of several vertebrates have challenged the paradigm that sIgA and sIgT are the only sIg required for microbiome homeostasis (*5*). For instance, in humans, sIgM is found in the gut mucus where it coats a significant proportion of gut microbiota along with sIgA (*5*).

This finding raised the notion that sIgM could help sIgA with shaping and retaining the gut microbiome (*5*). Similarly, we have previously reported that gut microbiota in rainbow trout is not only coated by sIgT, but also by sIgM (*6*). Interestingly, laboratory mice appear to show no sIgM coating of gut microbiota (*5, 7*), thus precluding the use of this model system for investigating the specific contribution of sIgM in the maintenance of microbiota homeostasis. Due in part to the latter, the functional significance of sIgM coating of gut microbiota across vertebrates remains a mystery.

Coating of microbiota by mammalian sIgA and fish sIgT appears to be fundamental for two paradoxical but conserved biological roles. On the one hand, microbiota coating by these two Igs promotes colonization of beneficial microbes at vertebrate mucosal surfaces (*4, 8*). On the other hand, sIgA/sIgT coating of pathogens or pathobionts prevents detrimental bacteria from penetrating the host mucosal barrier (*4, 9*). Additionally, both sIgA and sIgT are required for control of mucosal pathogens (*4, 8*). Interestingly, IgT is so far the most ancient Ig isotype specialized in mucosal immunity (*6, 10*). Yet, IgT is not found in several basal jawed vertebrate taxa including cartilaginous fishes or Holostei (bowfishes and gars). Moreover, IgT has been lost in several branches of Teleostei including Siluriforms, Beloniforms, Gobiforms and others (*4, 11*). This phylogeny suggests that other sIgs perform microbiota coating functions. Because of the ancient and functionally conserved nature of IgM in all jawed vertebrates and the recently reported conserved coating of fish and human gut microbiota by this Ig (*4, 5*), we hypothesized that sIgM plays a previously unanticipated role in shaping the gut microbiome and maintaining gut homeostasis. Using a novel *in vivo* model for IgM depletion in an early vertebrate, the teleost fish rainbow trout, we show that sIgM is required to maintain microbiota homeostasis in the trout gut. Lack of sIgM causes severe gut barrier damage, dysbiosis, inflammation, bacterial translocation and weight loss. Interestingly, changes in the microbiome of IgM-depleted animals are predictive of significant shifts in several important bacterial metabolic pathways known to be critical for host metabolism. Furthermore, trout devoid of sIgM are more susceptible to DSS-induced colitis thus implying a protective role of sIgM against the deleterious effects of colitis damage.

Besides unveiling a new role for sIgM in mucosal immunity, our work reveals that the interactions between sIgM and microbiota are not only the most ancient and conserved sIg-microbiota interactions in jawed vertebrates but are also required for the preservation of microbiota homeostasis, and suggests that metabolic mutualism is a major driver for this co-evolutionary process.

### Generation of IgM-depleted fish

An anti-trout IgM monoclonal antibody (mAb) (*12*) was used here to deplete IgM^+^ B cells *in vivo* by delivery of the mAb through intraperitoneal (IP) injection (fig. S1A). We initially used several anti-IgM mAb doses to assess depletion levels of blood IgM^+^ B cells after 6 weeks post injection in 2-3 g fish. We found that the optimal dose to achieve near complete depletion of blood IgM^+^ B cells was that of 300 μg of anti-IgM mAb/fish (fig. S1C) while a lower dose (200 μg/fish) led to a significant, but incomplete depletion of blood IgM^+^ B cells (fig. S1B). In contrast, IgM^+^ B cells were not depleted in fish treated with equal amounts of isotype mouse IgG control antibody (fig. S1, B and C). Importantly, none of the anti-IgM mAb doses or the respective isotype mouse IgG control Ab treatments induced any depletion of blood IgT^+^ B cells, thus supporting the specificity of the IgM^+^ B cell depletion treatment (fig. S1, B and C). Based on the aforementioned data, for the in vivo studies performed thereafter we used the dose of 300 μg/fish of anti-IgM mAb or its respective isotype control Ab dose.

To evaluate the degree and duration of IgM^+^ B cell depletion in both the blood and the gut, as well as its effect on serum IgM and gut mucus sIgM levels, we performed a time course study. When compared to control levels, IgM^+^ B cells were depleted almost completely both in blood and gut (69.5-98.9%) from week 1 up until week 9 post-depletion treatment (Fig. 1, A and B, left panels). At week 13 post depletion, IgM^+^ B cell numbers had recovered to the levels of control fish (Fig. 1, A and B, left panels). Interestingly in IgM-depleted animals we observed a considerable compensatory response in the % of blood IgT^+^ B cells which remained significantly higher than those of control fish throughout the treatment (Fig. 1, A and B, right panels). This IgT^+^ B cell-mediated compensatory response was observed in the gut of IgM-depleted fish only at weeks 4 and 6 post-depletion treatment (Fig. 1, A and B, right panels). The depletion treatment also led to the depletion of serum and gut mucus IgM (Fig. 1C) and sIgM (Fig. 1C) respectively. More specifically, depletion of serum IgM followed a similar trend to that IgM^+^ B cells while in the gut mucus, sIgM was not significantly reduced until week 3 post-depletion treatment (Fig. 1, A to D). Both in the serum and gut mucus the depletion treatment induced a near complete depletion of IgM and sIgM respectively at 6 and 9 weeks post-depletion treatment. IgM and sIgM concentrations recovered to those of control levels 13 weeks post-depletion treatment (Fig. 1, C and D). In contrast to IgM and sIgM, serum and gut mucus IgT and sIgT levels remained unchanged throughout the course of the experiment, thus showing a lack of IgT and sIgT compensatory responses (Fig. 1, E and F). In conclusion, we developed a strategy to selectively and consistently induce a near complete depletion of IgM^+^ B cells, IgM and sIgM in all tissues analyzed without depleting the levels of IgT^+^ B cells, IgT and sIgT.

**Fig. 1.**
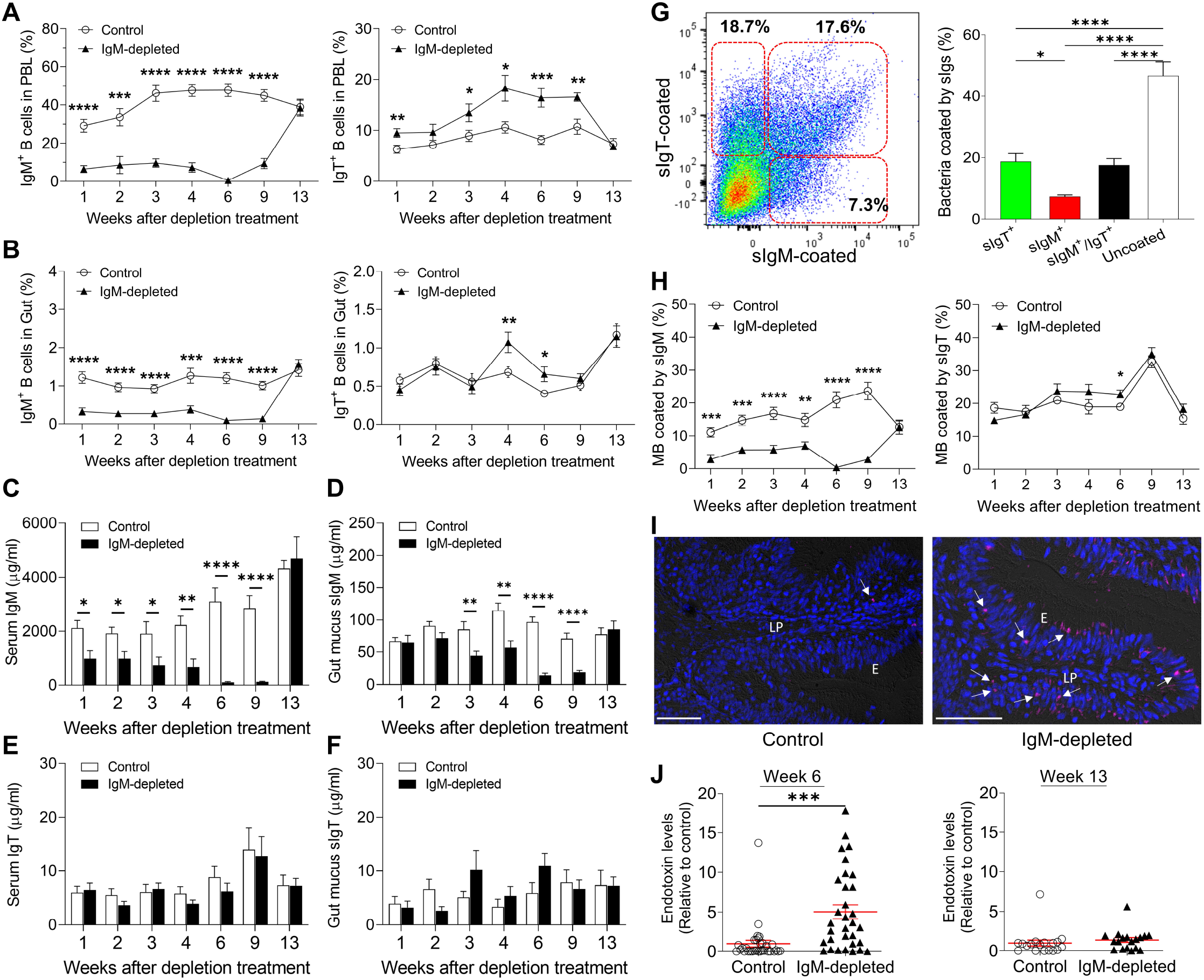
IgM depletion leads to drastic reductions of IgM^+^ B cells and sIgM in trout tissues and fluids respectively while inducing a major decrease in sIgM coating of the gut microbiota and their translocation into systemic circulation. Fish were injected with either 300 μg of anti-trout IgM mAb (IgM-depleted) or 300 μg of the isotype control antibody (control). Percentages of IgM^+^ and IgT^+^ B cells, and sIgs-coating of gut microbiota was determined by flow cytometry at the indicated weeks post IgM^+^ B cell depletion treatment. (**A** and **B**) Percentage of IgM^+^ (left panels) and IgT^+^ B cells (right panels) within the lymphocyte population of blood (A) and gut (B) leukocytes in control and IgM-depleted fish at the indicated weeks. Data are representative of at least two independent experiments (*n* = 13-14 fish per group, means ± SEM). (**C** to **F**) IgM and IgT concentration in serum and gut mucus upon IgM^+^ B cell depletion treatment. *n* = 12 fish per group. Data are representative of at least two independent experiments (means ± SEM). (**G**) Flow cytometry analysis of sIgM- and sIgT-microbiota coating from gut mucus of naïve fish. Representative dot plot (left) showing the proportions of sIgM- and sIgT-coated bacteria. Numbers adjacent to outlined areas indicate percentage of sIgT-coated microbiota (top left), sIgM-coated microbiota (bottom right) or sIgM/IgT-double coated microbiota (top right) in the bacterial population. Bar plot (right) showing the percentage of sIgM-or sIgT-coated microbiota from gut mucus of naïve fish (*n* = 8 fish per group). Data are representative of at least two independent experiments (means ± SEM). (**H**) Percentage of gut microbiota coated with sIgM (left) or sIgT (right) at the indicated weeks post-depletion treatment (*n* = 12 fish per group). Data are representative of at least two independent experiments (means ± SEM). (**I**) Detection of translocated bacteria by fluorescence *in situ* hybridization in gut cryosections from control (left) and IgM-depleted fish (right) at 6 weeks post IgM-depletion treatment. Gut cryosections were stained for Eubacteria detected with Cy5-EUB338 (magenta) and nuclei (blue) detection. Arrows indicate translocated intestinal bacteria through the epithelium and lamina propria. Scale bars, 50 μm. Abbreviations: E, epithelia; LP, lamina propria. (**J**) Endotoxin level measurements by LAL chromogenic endpoint assay in serum from control and IgM-depleted fish at 6 weeks (*n* = 32 fish per group) and 13 weeks (*n* = 18 fish per group) post-IgM depletion treatment. Sera endotoxin levels in IgM-depleted fish were normalized to those in control fish, which were set as 1. Each symbol represents an individual fish; small horizontal red lines indicate the means. Data are pooled from and representative of at least three independent experiments. Statistical analysis was performed by unpaired Student’s *t*-test (A to F, H and J) or by One-way ANOVA (G). \**P* < 0.05, \*\**P* < 0.01, \*\*\**P* < 0.001 and \*\*\*\**P* < 0.0001.

### Decreased sIgM-coating of microbiota correlates with increased levels of serum endotoxin

We next evaluated whether sIgM depletion had an impact on the gut microbiota coating by sIgM and sIgT. In Fig. 1G we show the proportions of sIgM and sIgT gut microbiota coating measured in naïve fish. We found that ∼18.7% and ∼7.3% of the microbiota were single-coated by sIgT and sIgM respectively while ∼17.6% were double-coated with sIgT and sIgM (Fig. 1G). Upon sIgM depletion, there was a drastic reduction (∼53.3-97.9%) in the % of sIgM-coated microbiota from week 1 until week 9 post-depletion treatment (Fig. 1H, left). In contrast, no significant changes in sIgT coating were observed at most time points except at week 6 for the same animals (Fig. 1H, right), thus indicating an overall lack of compensatory sIgT responses in microbiota-coating in the absence of sIgM. By week 13 post-depletion treatment, levels of microbiota coating by sIgM were fully recovered (Fig. 1H, left), thus mirroring the recovery of sIgM in gut mucus of IgM-depleted fish at the same time point (Fig. 1C). We then chose fish from the time point at which we observed the highest reduction in sIgM coating (week 6) to evaluate whether in the absence of sIgM coating, bacteria could still be contained within the luminal area or was able to translocate into the gut tissue. As shown in Fig. 1I, a considerable number of bacteria penetrated the gut tissue of IgM-depleted fish when compared to control animals. We then hypothesized that translocated bacteria might have entered systemic circulation. In support, we observed significant increases of LPS levels in the serum of a substantial number of IgM-depleted fish (Fig. 1J) at the same time point (week 6). In contrast, at week 13 post-depletion treatment, LPS levels of IgM-depleted fish returned to those of control animals, probably due to the recovery of sIgM microbiota coating at that time point (Fig. 1 H, left panel). Combined, these results strongly suggest a critical homeostatic role for sIgM in containing microbiota within the gut luminal area and in avoiding its systemic dissemination.

### IgM depletion induces severe gut tissue damage and inflammation

We next hypothesized that the translocation of microbiota into the gut tissue of IgM-depleted fish induced tissue damage and inflammation. Confirming this hypothesis, we observed acute changes in histopathology scores at 3 and 6 weeks post-depletion compared to controls (Fig. 2, A to C). Tissue damage was evidenced by severe lamina propria (LP) detachment from the epithelium, inflammation, edema, immune cell infiltrates and some degree of epithelial damage (Fig. 2, B and C). Supporting the inflammation detected at the histological level we also observed the upregulation of several pro-inflammatory cytokines mostly at 6 weeks post-depletion (Fig. 3D). More specifically, we found a ∼1.4-13.2 fold increase relative to control fish for *il-1β, tnfa½, il-17a/f2b, il-17a/f3, il-17a/f1b, il-21a*, and *il-22* transcripts. With regards to anti-inflammatory genes, *il-10a* and *il-10b* were up-regulated (∼2.9-3.2 fold increase) also at week 6 post-depletion (Fig. 3D), while *tgf1b* was mildly down-regulated (∼0.73-0.78 fold decrease), at weeks 1-6 post-depletion (Fig. 3E). Translocation and dysbiosis of microbiota at mammalian mucosal surfaces have frequently been correlated with increases in the gene expression of nucleotide binding oligomerization domain (NOD) and antimicrobial peptide (AMP) molecules (*13*). Similarly, at 3 and/or 6 weeks post-depletion we observed moderate increases (∼1.6-3.2 fold relative to control fish) in transcript levels of two LPS binding proteins (*lbp/bpi1* and *lbp/bpi2*), the AMP *cathelicidin* (*cath2*), and *lysozyme* (*lyz*) (Fig. 3F). A number of cytokines and AMP genes remained unchanged throughout the depletion period (fig. S2). Together, our results highlight the relevance of sIgM in the maintenance of mucosal barrier integrity, and suggest that in its absence, strong gut tissue inflammation occurs through translocation of dysregulated microbiota.

**Fig. 2.**
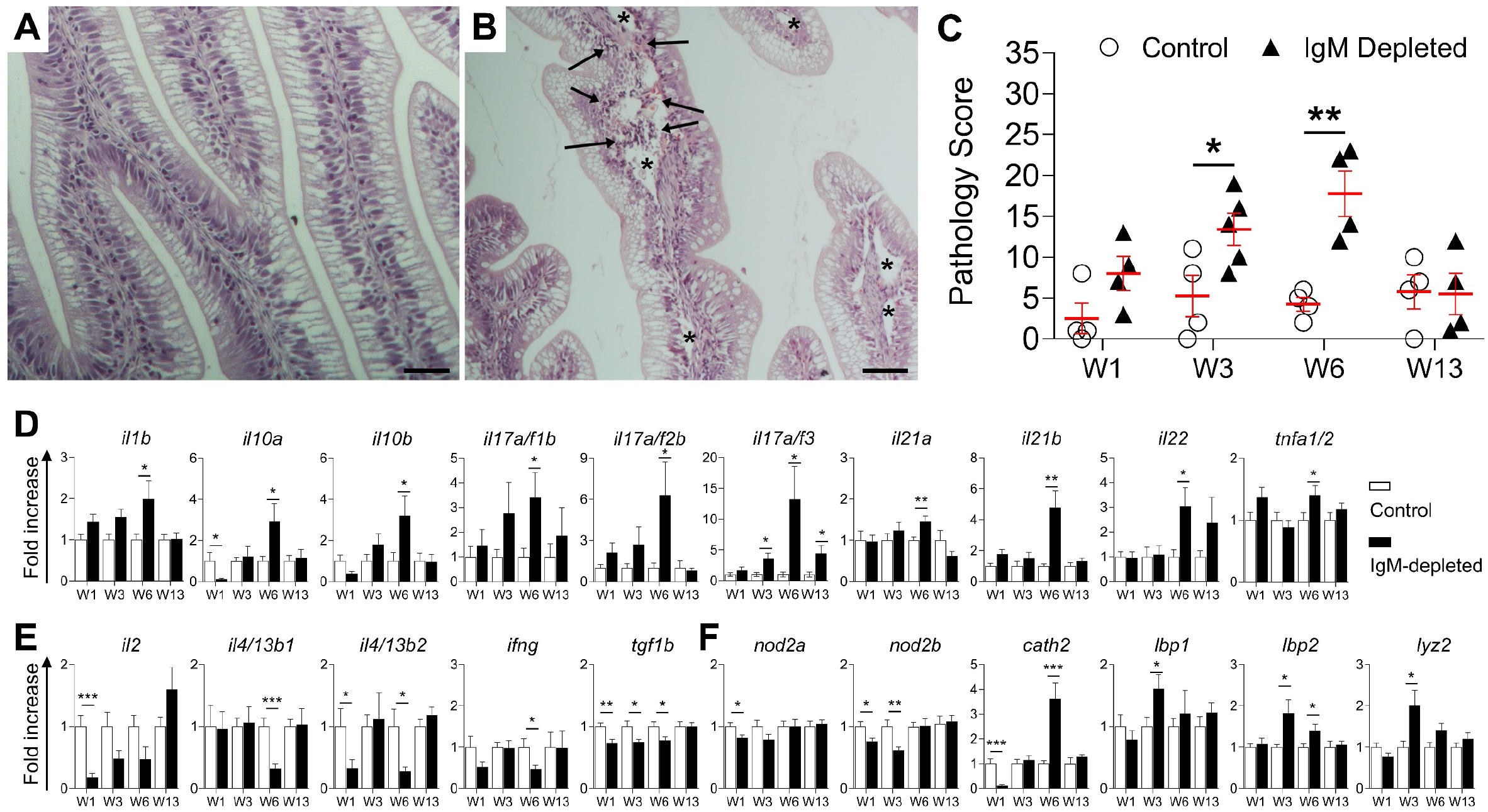
IgM depletion leads to severe tissue damage and inflammation. (**A** and **B**) Hematoxylin-eosin staining of gut paraffin sections in control (**A**) and IgM-depleted (**B**) fish at 6 weeks post-depletion treatment. Arrows indicate immune cell infiltrates, * indicate edema causing detachment of LP from epithelium. Scale bars, 50 μm. (**C**) Pathology score of gut tissue in control and IgM-depleted fish at the indicated weeks post IgM-depletion treatment (means ± SEM; *n* = 4-5 fish per group). (**D** to **E**) Real-time PCR analysis of cytokine genes that are up-regulated (**D**) or down-regulated (**E**), and antimicrobial peptide genes (**F**) in gut tissue from control and IgM-depleted fish at the indicated weeks post IgM-depletion treatment. Transcriptional levels in IgM-depleted fish were normalized to those in control fish, which were set as 1 (means ± SEM; *n* = 10 fish per group). Data are representative of at least two independent experiments. Statistical analysis was performed by unpaired Student’s *t*-test. \**P* < 0.05, \*\**P* < 0.01 and \*\*\**P* < 0.001.

**Fig. 3.**
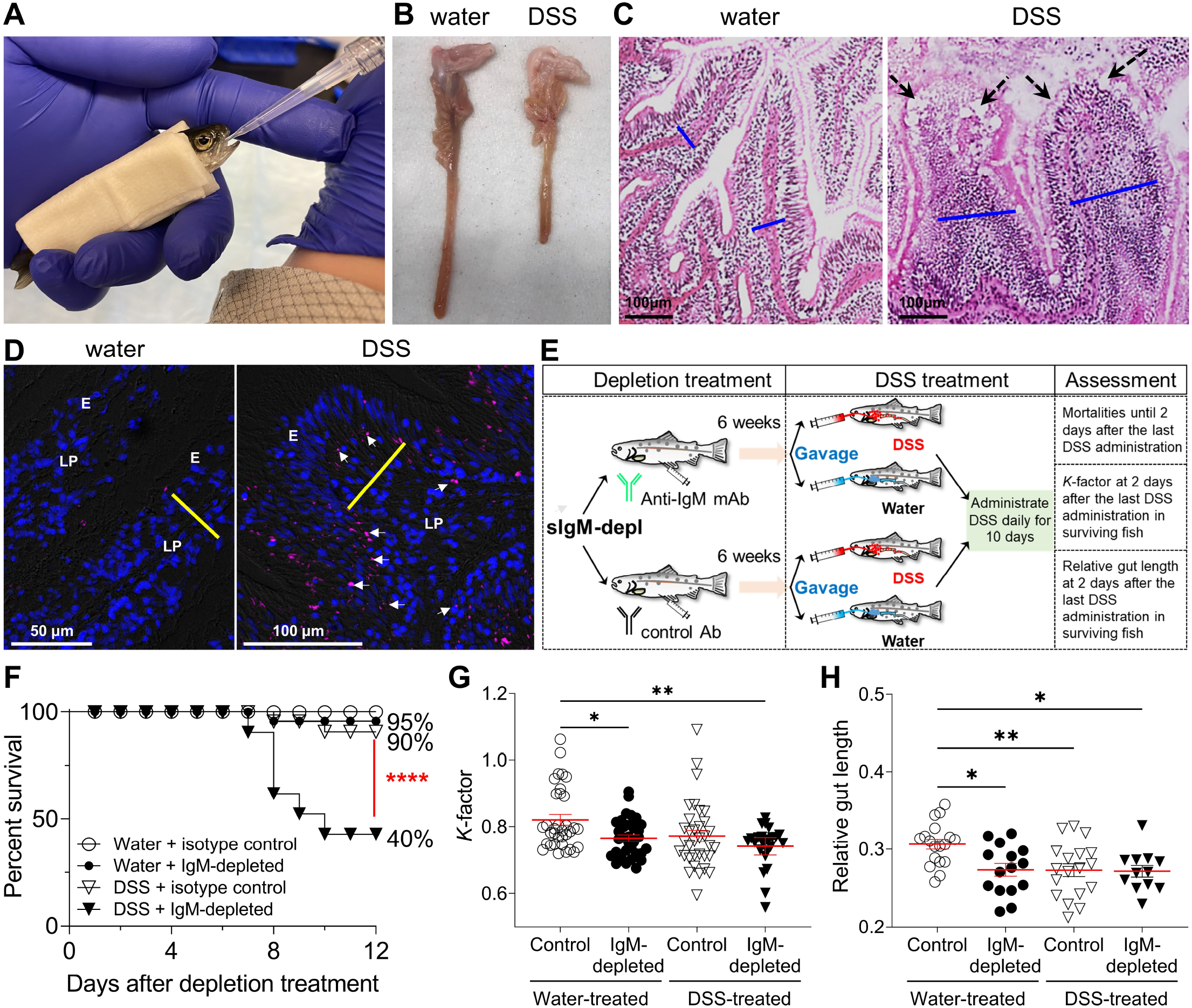
DSS-induced colitis induces significant fish mortalities in IgM-depleted fish. (**A**) Water or DSS (3% in water) administration (100 μl/fish) by oral gavage using a modified syringe attached to a pipette capillary tip (**B**) Gross morphology of the gastrointestinal tract of the water- and DSS-treated trout. (**C**) H&E staining images of the gut of water- and DSS-treated trout. Black dash-arrows point to epithelial damage, while the blue solid lines indicate the thickness of the lamina propria (LP). (**D**) Detection of bacteria by fluorescence *in situ* hybridization in gut cryosections from water-(left) and DSS-(right) treated trout. Gut cryosections were stained with Cy5-EUB338 for Eubacteria (magenta) detection and with DAPI for nuclei (blue) detection. Arrows indicate translocated intestinal bacteria through the epithelium and lamina propria. Abbreviations: E, epithelia; LP, lamina propria. (**E**) DSS-induced colitis in model IgM-depleted trout and control fish. Trout were injected with either anti-IgM mAb or isotype control Ab, and quarantined for 6 weeks. Thereafter fish were orally gavaged with a 3% DSS water solution or water alone, daily for 10 days. Fish mortalities were recorded for twelve days since day 1 of the DSS-treatment, while body weight, total length and gut length were measured from the surviving fish at the end of the experiment to calculate *K*-factor and relative gut length RGL. (**F**) Kaplan-Meier curves showing the percent survival rates following the water and DSS treatment in IgM-depleted and control fish. Numbers adjacent to symbols indicate percentage of survival rate. *n* = 20 fish per group. Data are representative of four independent experiments. (**G**) *K*-factor of the surviving fish following water and DSS treatment in IgM-depleted and control fish, *n* = 24-34 fish per group. Data are pooled from four independent experiments. (**H**) Relative gut length of the surviving fish following water and DSS treatment in IgM-depleted and control fish, *n* = 12-18 fish per group. Data are representative of four independent experiments. Statistical analysis was performed by Logrank (Mantel-Cox) test (F) or One-way ANOVA (G and H). \**P* <0.05, \*\**P* <0.01, and \*\*\*\**P* <0.0001.

### DSS-induced colitis exacerbates fish mortalities in IgM-depleted fish

The bacterial translocation, along with the tissue damage and inflammation induced in the gut tissue upon IgM-depletion suggests a protective role of sIgM against colitis-like pathology. To confirm this hypothesis, we induced colitis in sIgM-depleted fish with the goal to evaluate whether absence of sIgM would render IgM-depleted fish more susceptible to colitis-induced mortalities. To this end, we first developed a novel DSS-induced colitis trout model. To induce colitis, fish were gavaged with a solution of 3% DSS in water, daily for 10 days (see Fig. 3A). Control fish were gavaged with water. As seen in Fig. 3B, the intestines of DSS-treated fish were significantly shortened. Moreover, H&E staining of gut sections from DSS-treated fish showed significant epithelial and tissue damage as well as immune cell infiltration and thickening of the lamina propria (Fig. 3C). Importantly, we detected visible translocation of microbiota in the trout gut tissue of DSS-treated fish (Fig. 3D). When we induced DSS-colitis in sIgM-depleted animals (Fig. 3E), fish mortalities increased dramatically from 10% in DSS-control fish to 60% in DSS-IgM-depleted fish (Fig. 3F). Mortalities in the IgM-depleted fish gavaged with water were only 5% thus indicating that the mortalities in the DSS-IgM-depleted group were exacerbated by DSS treatment. Interestingly, when compared to control fish gavaged with water, the *K*-factor and relative gut length (RGL) were significantly reduced in the IgM-depleted fish gavaged with water, while these measurements in the later group were similar to the levels observed in the DSS-control group (Fig.3, G and H). This indicated that the effect of IgM-depletion induced similar reductions in body weight and RGL than that of DSS-induced colitis in control animals. However, these two parameters were not further reduced in IgM-depleted fish treated with DSS. In conclusion, these data suggest a previously unrecognized protective role of sIgM against the deleterious effects of colitis damage.

### IgM-Seq reveals overlapping microbiota coating patterns among sIg isotypes

Our previous work in trout gill revealed that sIgT coats a specific subset of bacterial taxa with both beneficial and pathobiont properties (*10*). The aforementioned study, however, did not investigate what bacteria are targeted by sIgM. Therefore here we sorted single sIgM^+^-, single sIgT^+^- and double sIgM/sIgT^++^-coated bacteria as well as uncoated bacteria from control trout gut mucus samples and performed 16S rDNA amplicon sequencing as previously described (*10*). Our results indicate that in terms of composition, microbial community targeted by sIgM is largely similar to that of the pre-sorted community as well as the sIgM/sIgT^++^ and the uncoated communities (Fig. 4, A to D). In support, alpha diversity metrics were not significantly different among the four sorted bacterial populations (Table S1). Principal component analysis revealed strong overlap between the presort, sIgM^+^ and sIgM/sIgT^++^ communities and more distinct separation of the sIgT^+^ and uncoated microbial communities with respect to the rest (Fig. 4A).

**Fig. 4.**
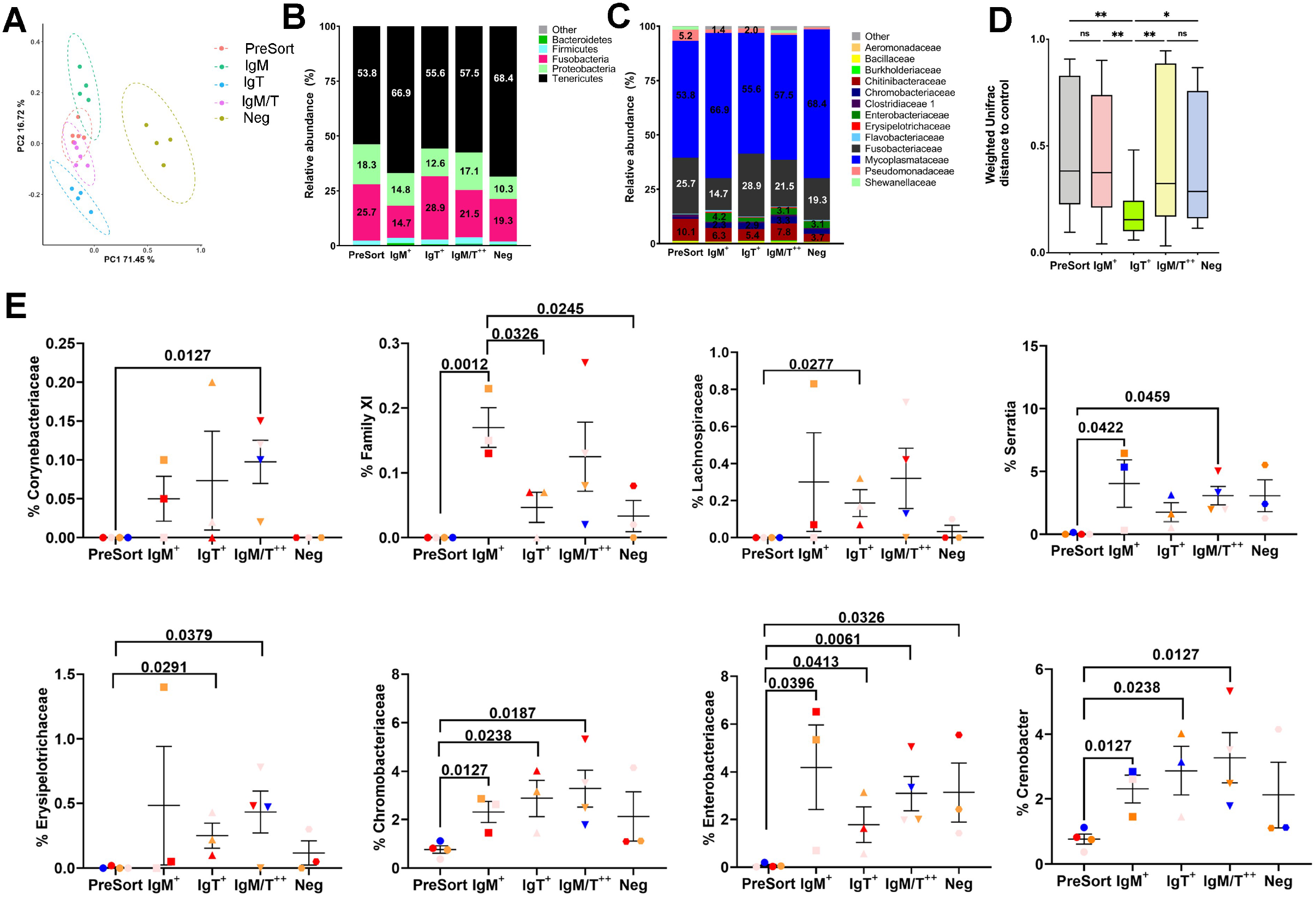
Ig-Seq reveals overlapping coating patterns by sIgM and sIgT in trout gut. Total bacteria and sIgM^+^-, sIgT^+^-, sIgM/sIgT^++^-coated and uncoated bacteria from the gut mucosa were flow cytometrically sorted and high-throughput 16S rDNA sequencing of the V1 to V3 region was performed (Ig-seq). (**A**) Principal coordinate analyses of the pre-sort, sIgM^+^, sIgT^+^, sIgM/sIgT^++^ and uncoated microbial communities from sorted gut bacteria obtained from control trout (*n=*3-4). (**B**) Relative abundance at the phylum level of pre-sort, sIgM^+^, sIgT^+^, sIgM/IgT^++^ and uncoated microbial communities in rainbow trout gut. (**C**) Relative abundance at the family level of pre-sort, sIgM^+^, sIgT^+^, sIgM/sIgT^++^ and uncoated bacterial communities in rainbow trout gut. (**D**) Weighted Unifrac distance of the pre-sort, sIgM^+^, sIgT^+^, sIgM/sIgT^++^ and uncoated microbial communities in rainbow trout gut. (**E**) Relative abundance of significantly different ASVs detected in the pre-sort, sIgM^+^, sIgT^+^, sIgM/sIgT^++^ and uncoated bacterial communities in rainbow trout gut. \**P* <0.05. \*\**P* <0.01.

Interestingly, the weighted unifrac distance of the sIgT^+^ microbial community was the only one which was significantly different to all the other sequenced communities, with no differences in the weighted unifrac distance between the sIgM-coated microbial community and the others (Fig. 4D). At the phylum level we noted some differences such as the higher percentage of Tenericutes of the family *Mycoplasmataceae* represented in the sIgM^+^ coated community (Fig. 4, B and C) albeit this was not significantly different. We observed significant preferential coating of *Enterobacteriaceae* such as the potential pathogen *Serratia sp*. and *Family XI* (*Clostridiales*) by sIgM compared to the other groups (Fig. 4E). Two key short chain fatty acid (SCFA) producers, *Lachnospiraceae* and *Erysipelotrichaceae*, were preferentially coated by sIgT whereas *Corynebacteriaceae* abundance was significantly higher in the double coated community (Fig. 4E). We did not detect amplicon sequence variants (ASVs) that were uniquely coated by sIgM only or sIgT only indicating overlapping coating patterns by both sIg isotypes in the trout gut. Combined, our results indicate preferential coating of sIgM of some taxa such (i.e, *Enterobacteriaceae, Family XI*) but a clear convergent coating pattern of microbiota by sIgM and sIgT.

### IgM depletion causes gut dysbiosis

Given the tissue damage, inflammation and endotoxin levels in circulation in IgM-depleted animals, we next aimed to determine how absence of sIgM perturbs the microbial community composition of the trout gut. IgM depletion caused significant losses in the gut mucus-associated bacterial loads 6 weeks post-depletion as measured by qPCR (Fig. 5A). Microbiome profiling by 16S rDNA amplicon sequencing identified a significantly higher Shannon Diversity Index 1 week post-depletion in IgM-depleted compared to control trout gut suggesting colonization of novel taxa at this time point (Fig. 5B). No other significant differences in other alpha diversity metrics were observed (Table S2). IgM depletion resulted in significant changes of the gut microbial community composition. The weighted unifrac distance of the IgM-depleted gut microbial community 1 week post-depletion was also significantly different from its corresponding controls, indicating quick shifts in the gut microbial community following IgM depletion treatment (Fig. 5C). At the phylum level, IgM depletion reduced the proportion of Tenericutes by week 1 at the expense of expansions of Proteobacteria, Firmicutes and Fusobacteria (Fig. 5D). The decreased Tenericutes abundance was due to the decrease of the family *Mycoplasmataceae* (Fig. 5E). At the genus level, IgM depletion resulted in a sharp decrease in *Mycoplasma sp*. abundance (Fig. 5F) and an increase in *Deefgea sp*. and *Cetobacterium sp*. abundances at week 1. The abundance of *Mycoplasma sp*. was also significantly lower in IgM-depleted fish at weeks 6 and 13 (Fig. 5F) and increased *Deefgea sp*. levels were still detected at week 13, indicating some long-lasting changes in microbial community composition even when IgM levels are restored (week 13). Interestingly, *Clostridium sp*. abundance, a producer of SCFA, increased at week 1 and decreased at week 3 compared to controls. We also detected significant changes in the abundance of three other ASVs over time, *Holosporaceae, Lachnospiraceae* and *Shewanella sp*, a pathogen of fish (Fig. 5F). Interestingly, *Lachnospiraceae* was noted as one of the taxons preferentially coated by sIgT in the trout gut, suggesting that shifts in response to sIgM depletion can also indirectly impact taxa preferentially coated by other sIg isotype. Of note, *Lachnospiraceae* abundance was significantly lower at weeks 3 and 13 in IgM-depleted compared to the control group.

**Fig. 5.**
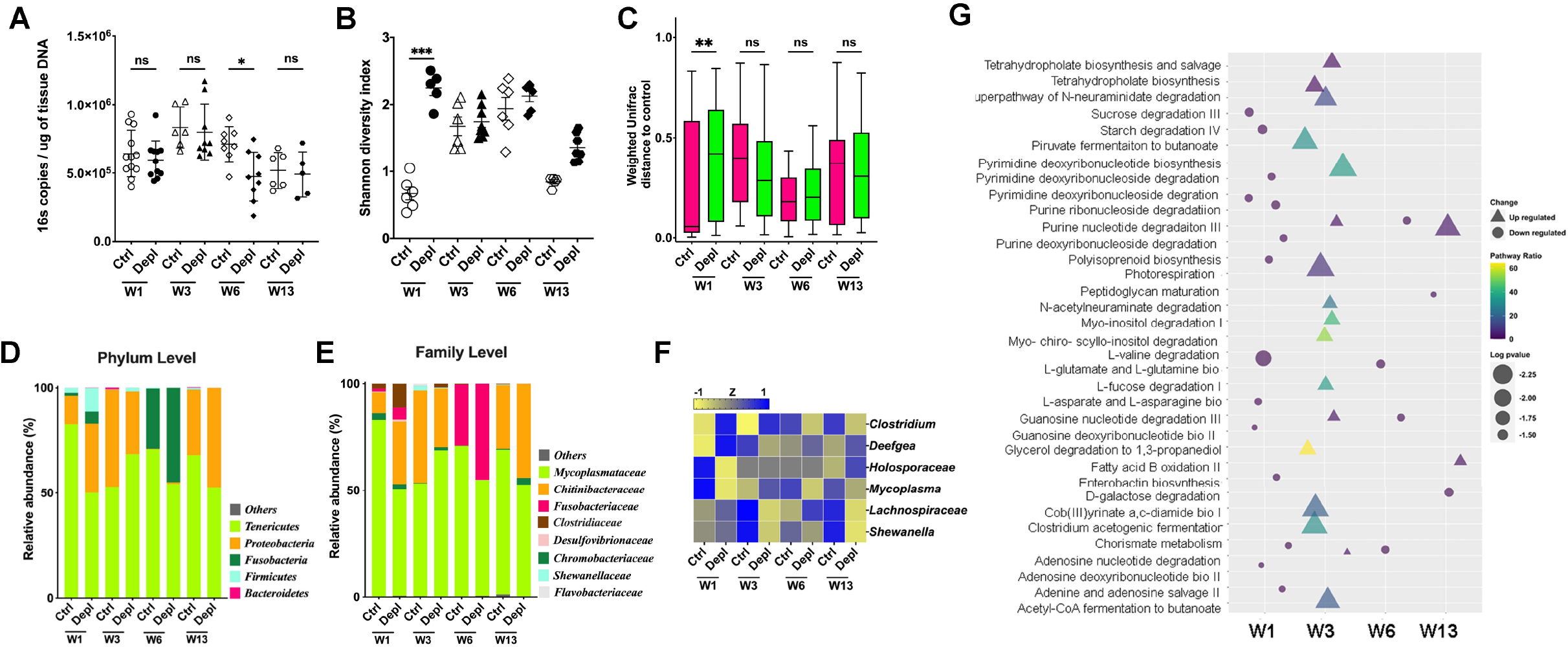
IgM depletion leads to gut microbiota dysbiosis. The microbial community composition of control and IgM-depleted gut was determined by high-throughput 16*S* rDNA sequencing at 1, 3, 6 and 13 weeks post-depletion treatment. **(A)** Bacterial loads in the gut of control and IgM-depleted trout 1, 3, 6 and 13 weeks post-depletion as measured by qPCR (*n* = 9 fish per group). (**B**) Mean Shannon Diversity Index of the gut microbial community of control and IgM-depleted trout 1, 3, 6 and 13 weeks post-depletion (*n* = 9 fish per group). (**C**) Weighted Unifrac distance of control and IgM-depleted gut microbial communities 1, 3, 6 and 13 weeks post-depletion (*n* = 9 fish per group). (**D**) Relative abundance at the phylum level of the gut microbial community composition of control or IgM-depleted trout 1, 3, 6 and 13 weeks post-depletion (*n* = 9 fish per group). (**E**) Relative abundance at the family level of the gut microbial community composition of control or IgM-depleted trout 1, 3, 6 and 13 weeks post-depletion (*n* = 9 fish per group). (F) Heat map with the significantly different ASVs in control and IgM-depleted gut microbial communities 1, 3, 6 and 13 weeks post-depletion (*n* = 9 fish per group). (G) PICRUSt2 analysis of the microbial communities of control and IgM-depleted gut microbial communities 1, 3, 6 and 13 weeks post-depletion showing the predicted downregulated and upregulated biological pathways in each community. \**P* <0.05.

Analysis of predicted biological pathways encoded by the bacterial communities of each treatment group at the four time points using PICRUSt2 revealed striking findings (Fig. 5G). At week 1, all significant metabolic pathways were down-regulated in IgM-depleted compared to control fish, the most down regulated pathway was L-valine degradation. The peak of the predicted pathway changes occurred at week 3, when all significantly different pathways were upregulated in IgM-depleted fish. Upregulated pathways included several amino acid metabolism pathways, pyrimidine and purine biosynthesis and degradation pathways and carbon metabolic pathways such as pyruvate fermentation to butanoate (butyrate) or acetyl-coA fermentation to butanoate (butyrate) (Fig. 5G). PICRUSt2 detected four significantly regulated pathways at weeks 6 and 13, suggesting that IgM depletion causes long-term perturbations in the predicted metabolic capacities of the trout gut microbial community. Combined, these results indicate that IgM is not only necessary for maintenance of microbial community composition in the gut, but suggest also that IgM coating of microbiota is involved in the metabolic relationship between host and microbiota.

## Discussion

The co-evolution between microbiota and the host adaptive immune system has created a conserved and mutualistic relationship in all jawed vertebrates (*14*). Over 500 million years ago, IgM first emerged in cartilaginous fish, and homologs of this ancient IgM has been identified in every jawed vertebrate taxon. Thus, the phylogeny of IgM clearly indicates a striking, high degree of conservation of this Ig isotype since the inception of B and T cell-based adaptive immunity (*1, 2*). Both in cold- and warm-blooded vertebrates, IgM has always been viewed as a systemic immunoglobulin that plays a key role against infection, while its role in mucosal immunity and microbiota regulation has been vastly overlooked (*2, 4*).

Until recently it was believed that mammalian sIgA was the only sIg involved in the maintenance of gut microbiota homeostasis (*15-17*). Our follow-up studies on fish sIgT demonstrated that control of microbiota homeostasis by sIgs is a primordial function of specialized mucosal sIgs already found in early vertebrates such as teleosts (*10*). However, the current dogma that sIg-dependent microbiota homeostasis is mediated by sIgA or sIgT in vertebrates, has recently been challenged by several studies showing that significant proportions of human and fish microbiota are coated by sIgM in the gut and other mucosal surfaces (*5, 18, 19*). Because sIgM does not coat the microbiota of mice under either specific-pathogen-free (SPF) or non-SPF conditions (*5, 7*), laboratory mouse models to study the potential role of sIgM in host-microbiome interactions are not suitable to address this question. Thus, the question remains as to whether sIgM plays a novel role in mucosal immunity, and thus, is an Ig required in the maintenance of microbiome homeostasis.

We therefore developed a strategy to selectively deplete IgM^+^ B cells in rainbow trout, a teleost species. This strategy was highly effective in depleting most of the IgM and IgM^+^ B cells from all tested fluids and lymphoid organs respectively, for over a 9-week period. We have previously used depleting mAbs to deplete sIgT and IgT^+^ B cells and others have shown in mammalian systems that B cell depletions covering several weeks work well with the use of a single antibody treatment (*20, 21*). To our knowledge, our IgM depletion model represents the only vertebrate in vivo model that allows for the temporal and selective depletion of gut sIgM, thus enabling the evaluation of the role of sIgM in microbiome homeostasis. Here we show that a large portion of the trout gut microbiota is double-coated by sIgT and sIgM, while the largest proportion is sIgT single coated, being the smallest percentage single sIgM coated. Interestingly, two reports have shown that the human gut microbiota follows a similar pattern of coating with sIgA and sIgM double-coated microbiota being the most prevalent coating mode (*5, 18*), while in another human study they reported different proportions of sIgA/sIgM coating (*19*). Among many other possibilities, it is likely that such differences in gut sIg microbiota coating in humans are due to methodological methods in the assessment of sIg coating, or the use of different human cohorts with very different genetic backgrounds.

Depletion of sIgM led to a striking decrease in the sIgM coating of the trout gut microbiota that lasted for a 9 week period. Interestingly, we did not see a compensatory sIgT coating response in the IgM-depleted fish, in contrast with the compensatory sIgM coating response that occurs in the gill mucosa upon depletion of sIgT (*10*). This lack of compensatory sIgT coating may explain the severe phenotypes observed in the gut of sIgM depleted animals with regards to their tissue damage and inflammation. In addition to losing most of its sIgM coating, bacterial loads associated with the gut were reduced and large numbers of microbiota translocated within the gut tissue and significant amounts of LPS were found in the sera, thus indicating that sIgM is required for microbiota colonization and containment of sIgM-coated microbiota within the luminal area of the gut, as suggested also for sIgT and sIgA in fish and mammalian mucosal surfaces respectively (*10, 22*).

The significant gut tissue and systemic bacterial translocation observed upon sIgM depletion was accompanied with the occurrence of striking gut tissue damage and transcriptional upregulation of critical pro-inflammatory cytokines and AMPs. These data strongly suggest that the translocation of microbiota into the gut tissue leads to local inflammatory and antimicrobial responses similar to what has also been described in fish and murine models lacking sIgT and sIgA respectively (*10, 22*). It is interesting that sIgM depletion led to transcript increases of most of the gut cytokines also found to be upregulated in the gill of sIgT-depleted animals, albeit the increases in transcript levels were more marked upon depletion of sIgM (*10*), consistent also with the more severe tissue damage observed in sIgM-depleted animals in comparison to that of sIgT-depleted fish. Another meaningful difference not seen in the IgT-depleted fish (*10*) are the significant increases of *il-10* and *il-22* transcripts observed here at week 6 post-IgM depletion, when gut tissue damage was at its highest level. As these two cytokines play a key role in the maintenance of gut barrier function after injury, and stimulate the production of AMPs (*23, 24*), our findings suggest that their upregulation is induced in response to the more marked tissue damage and presence of translocated microbiota detected in the gut of IgM-depleted fish. It is worth pointing out that the aforementioned differences in cytokine transcript levels between IgT- and IgM-depleted fish maybe due in part to the different tissues examined as well as to the large differences in microbiota composition between the gill and the gut (*25*).

DSS-induced colitis in mice has become an indispensable model to investigate underlying mechanisms of colitis and understanding the roles of sIgA in the modulation of microbiota during dysbiosis (*7, 26-28*). Since lack of sIgM induces extensive tissue damage, inflammation and microbial dysbiosis, this implies that presence of sIgM is protective against the induction of colitis adverse effects. To confirm this hypothesis, we induced colitis in sIgM-depleted fish with the goal of evaluating whether absence of sIgM would render IgM-depleted fish more susceptible to colitis-induced adverse effects. To this end, we first developed a novel DSS-induced colitis model in fish in which the DSS solution was orally administered by gavage. Of note, previous DSS-induced colitis models have been developed in zebrafish, in which the whole larvae are immersed in a DSS-containing solution (*29, 30*). Our gavage strategy avoids potential side effects from exposing the whole organism in the DSS solution while providing a phenotype very similar to that observed in mice fed with DSS-water (*27*). As expected, upon induction of DSS-colitis in sIgM-depleted fish, mortalities increased considerably when compared to those of sIgM-depleted animals gavaged with water, thus providing further evidence of the protective role of sIgM against the deleterious effects of colitis damage. At this point we are uncertain of the specific mechanisms as to how sIgM exerts its protective effects in the gut. However, it is tempting to hypothesize that in the absence of sIgM, gut pathobionts are greatly expanded, as observed here (i.e., *Deefgea* and *Shewanella sp*, both of which can be pathogens (*31, 32*) of fish) and upon DSS-treatment, the severe gut epithelial damage induced enables further the translocation the expanded pathobionts into circulation, possibly causing sepsis and death.

Alternatively, it is possible that systemic IgM is important in the elimination of systemically escaped pathobionts, and thus, in the absence of serum IgM, bacteria cannot be eliminated, thereby causing sepsis and mortalities in fish. In that regard, it has been shown that microbiota can induce protective systemic IgA and IgG against polymicrobial sepsis or *E. coli* translocated from the gut respectively (*27, 33*). Future work is warranted to analyze these exciting possibilities.

While the microbial community coated by sIgA has been sequenced using IgA-Seq in several studies both in humans and mice (*5, 34*), the bacterial taxa targeted by sIgM has only been investigated in humans and to a much lesser degree when compared to that of sIgA. A recent study in humans found that the sIgM-coated community is encompassed by the sIgA-coated microbial community (*18*), a finding that suggests convergence of microbiota responses by sIgM and sIgA (*34*). In a different study, sequencing of sIgA^+^, sIgA/sIgM^++^ and uncoated human ileal and colonic bacterial populations found that double sIgA/sIgM^++^ coated bacteria proportionally had more Firmicutes but less Bacteroidetes (*5*). Our IgM-Seq experiments in trout gut did not find any taxa that were exclusively coated by sIgM or sIgT. However, our results indicate some degree of preferential coating by one or the other isotype as evidenced by the greater proportion of *Enterobacteriaceae* and *Family XI* in the single sIgM-coated community. Interestingly, the sIgT-coated bacterial community had a more defined beta diversity that was significantly different from the other groups (sIgM-coated, double coated and uncoated) suggesting that specialized sIgs may be coating certain bacterial taxa or that there is a dynamic continuum of coating and uncoating for sIgs that causes the results we obtained. Such changes in coating within the same bacterial species could be driven by changes in bacterial surface antigens during growth or by interaction of that species with other microbial species, as see in (*27, 33*).

IgM-depletion led to gut microbial dysbiosis, as previously observed also for IgT-depleted fish in the gill mucosa (*10*). In that regard, several conclusions can be made from the impact of IgM depletion on the trout gut microbiome. First, microbiota colonization in the gut mucosa requires IgM since bacterial loads in IgM-depleted trout were much lower than in control animals at the peak of the depletion. Interestingly, sIgT also promotes bacterial colonization in the trout gill but (*10*), but in the absence of sIgM, sIgT was not sufficient to maintain bacterial colonization levels in the trout gut. Second, certain taxa expand in the absence of sIgM control, such as *Deefgea sp*. and *Clostridium sp*. Interestingly, sIgT depletion also resulted in expansion in *Deefgea sp*. in the trout gill, again highlighting how specific microbiota taxa are particularly susceptible to host immune control mechanisms. Finally, absence of IgM led to long-lasting changes in the gut microbial community that appeared largely due to the decreased abundance at week 13 post depletion of taxa such as *Lachnospiraceae, Mycoplasma* sp. and *Serratia sp*. This indicates that some of the perturbation caused by sIgM depletion persists beyond the recovery of sIgM levels in the system and suggests long lasting detrimental effects on host gut physiology and metabolism.

The reciprocal interactions between microbiota and host immune systems are thought to be based on the fundamental metabolic benefits drawn by both partners through this interaction (*14, 35, 36*). It is logical to postulate, therefore, that the original relationship between microbiota and sIgM brought several evolutionary advantages to this partnership, likely through enhanced metabolic networks. In support, depletion of sIgM resulted in significant weight losses in trout, which in part could be due to the observed absence of beneficial microbiota colonizing the gut in the absence of sIgM. In that vein, studies in humans found that obese type 2 diabetes young individuals have more IgM-coated gut bacteria compared to age matched healthy controls and that upon fecal microbiome transplant into germ free B6 mice, recipient mice had impaired glucose tolerance and gained weight (*37*). The latter study supports the concept that lack of sIgM-microbiota coating promotes weight loss as indicated by our studies. We also noted that the dysbiotic gut microbiome resulting from depletion of sIgM is predicted to have very different metabolic capabilities compared to that of non-depleted controls. The predicted metabolic changes in the microbiome of IgM-depleted animals occurred early and included diverse pathways from amino acid and purine metabolism to carbon metabolism and production of SCFA. For example, because *Lachnospiraceae* (Order *Clostridiales*) are a family of obligate anaerobes that ferment diverse polysaccharides to SCFA and alcohols (*38*), the observed decreases of this family in IgM-depleted fish may result in altered SCFAs levels in the host. Interestingly, as previously reported in our sIgT-seq study in the trout gill (*10*), microbiota preferentially coated by sIgT in the gut have predicted roles in producing key metabolites such as SCFA. Our current efforts to perform metagenomics and metatranscriptomic analyses of each of the sIgT and or sIgM-coated bacterial communities should clarify further potential associations of sIg coating and specific microbial metabolic pathways.

In conclusion, the present study demonstrates that the most ancient and conserved immunoglobulin isotype, IgM, is essential for the maintenance of microbiota homeostasis while it also challenges the current paradigm that sIgA and sIgT are the key vertebrate sIgs implicated in that role. Moreover our findings suggest that evolution of mutualism between jawed hosts and their microbiota has been driven in part by the potential of sIgM in regulating microbial metabolism.

## Acknowledgments

We would like to thank D. Dinwiddie for sharing the Illumina sequencer; L. Bu for help with microbiome bioinformatics analysis; J. S. Faust and J. Fundyga of the Flow Cytometry Facility (Wistar Institute) for all the cell-sorting procedures conducted; The staff from the University Laboratory Animal Resources (ULAR) from the University of Pennsylvania for its daily effort displayed in the daily husbandry of our animal facilities and specially to Dr. A. Carthy for the excellent veterinary care of the rainbow trouts.

## Funding

National Institutes of Health 2R01GM085207-09 (JOS, IS)

U.S. Department of Agriculture grant USDA-NIFA 2021-67015 (JOS)

U.S. Department of Agriculture grant USDA-NIFA 2022-67015 (JOS) National Natural Science Foundation of China 31802337 (YD)

International Postdoctoral Exchange Fellowship Program by the Office of China Postdoctoral Council PC2018016 (YD)

Japan Society for the Promotion of Science Overseas Research Fellowship (YS) Japan Society for the Promotion of Science KAKENHI 20K22594 (YS)

Japan Society for the Promotion of Science KAKENHI 21H02288 (YS, FT)

## Author contributions

Conceptualization: JOS, IS

Methodology: YD, AFM, AM, EC, YS, FT, RM, WK, YY

Visualization: YD, AFM, AM, EC, YS

Funding acquisition: JOS, IS

Project administration: JOS, IS

Supervision: JOS, IS

Writing – original draft: JOS, IS

Writing – review & editing: JOS, IS, YD, AFM, AM, RM

## Data and materials availability

All data are available in the main text or the supplementary materials. The 16S rRNA sequencing data were deposited at the NCBI Sequence Read Archive (SRA) with accession numbers of Bioproject PRJNA938924 (for sIgM-seq) and PRJNA938926 (for gut microbiome of IgM-depleted fish).

## Supplementary Materials

Materials and Methods Figs. S1 to S2

Tables S1 to S3

References (*39–52*)

## Supplementary Materials for

Materials and Methods

Figs. S1 to S2

Tables S1 to S3

References

## Materials and Methods

### Fish maintenance

Juvenile rainbow trout (*Oncorhynchus mykiss*) from 2 to 3 g were obtained from Troutlodge and maintained as previously described (*1*). Briefly, fish were at least acclimatized for 2 weeks prior to start the experiments, in a RAS (recirculating aquaculture system) with water temperature of 15 ±1 °C and photoperiod of 10 hours per day. Fish were fed at 1% of biomass per day on a commercial pellet diet (Bio-Oregon). The selected rainbow trout are from the May strain and are not genetically inbred. This particular strain is an outbred strain that has been maintained as a closed population for at least nine generations. All animal procedures were approved by the Institutional Animal Care and Use Committees of the University of Pennsylvania. Depletion of IgM^+^ B cells and mucosal IgM (sIgM) from tissues and body fluids, respectively.

The in vivo depletion model has been adapted from the previously used strategy for IgT depletion (*10*). In order to initially evaluate the *in vivo* IgM^+^ B cell depletion effect of anti-trout IgM mAb, fish (∼2 to 3 g) were intraperitoneally injected with two different doses (200 or 300 µg) of mouse anti-trout IgM mAb (clone 1.14; IgG1) (*12*) or mouse IgG as control Ab (Innovative research). Fish were euthanized at three and six weeks upon Abs treatment and peripheral blood leukocytes (PBL) were obtained. The IgM^+^ and IgT^+^ B cells in PBL were stained and analyzed by flow cytometry as described below. The treatment of 300 µg of anti-trout IgM mAb was confirmed to successfully deplete IgM^+^ B cells over six weeks, and was selected to be used as the amount for the rest of IgM depletion procedure to achieve a consistent and high level of IgM^+^ B cell depletion in all fish. A dynamic time course of IgM^+^ B cell depletion from PBL and gut was thereafter performed using 300 µg per fish of anti-trout IgM mAb or its corresponding isotype control and evaluated by flow cytometry. Concentration of sIgT and sIgM in the serum and gut mucus of these fish were measured by western blot as described below. The biometrics including whole length in centimeters and body weight in grams were recorded in order to determine growth parameters. Fulton condition factor was calculated as *K*= 100*weight/length^3 (*39*).

### Collection of plasma, gut mucus and gut bacteria

Juvenile rainbow trout were euthanized by anesthetic overdose (MS-222, Syndel), and once fish dead was confirmed, the base of the tail was sectioned with a scalpel and blood was recovered with a heparinized capillary tube (*6, 40, 41*) Capillary tubes were centrifuged at 600 × g for 10 minutes at 4 °C, obtaining the buffy coats (i.e., leukocytes) and plasma. In order to collect gut mucus and the bacteria associated, fish were perfused with PBS-heparin (Sigma-Aldrich) through the heart to remove the circulating blood from the vessels associated to the gut, and confirmed with the gill arches were completely blanched. Therefore, the gut was extracted and opened longitudinally, and the surface was immersed in 200 µl of ice-cooled gut buffer [1× protease inhibitor cocktail (Roche), 0.2 mg/mL trypsin inhibitor from soybean (Thermo fisher Scientific), 1 mM PMSF (Sigma-Aldrich), 10 mM EDTA (Invitrogen), 0.5% bovine serum albumin (BSA, Gold Biotechnology) in PBS (PBS-BSA), pH7.2] as previously described (*6*).

The gut mucus was collected by gently scraping the inner surface of the gut, and transferred to a 1.75 ml Eppendorf tube. The sample was vigorously vortexed immediately to resuspend gut mucus with protease inhibiting gut buffer, and thereafter homogenized by passing through a syringe (needle 23G) 20 times. The samples were then centrifuged at 40 × g for 10 min to remove large particles, followed by 300 × g for 10 minutes at 4 °C three times in order to remove cell debris. The cell-free supernatant (containing gut mucus and bacteria) was centrifuged at 13,000 × g for 5 minutes at 4 °C to separate gut bacteria (pellet) from gut mucus (supernatant). Isolated bacteria were washed 3 times with 1% PBS-BSA (pH7.2) and finally resuspended in the same buffer for sIg-coating analysis. Gut mucus was immediately aliquoted and frozen at -80 °C until further use.

### Isolation of blood and gut leukocytes

Gut tissues were collected and pressed through a 100-μm cell strainer (BD Biosciences), and the leukocytes were suspended in an iced-cooled 1 × DMEM [Dulbecco’s modified eagle medium (DMEM, Thermo Fisher Scientific) supplemented with 1% FBS (Serum Source International) and 1% penicillin-streptomycin (Thermo Fisher Scientific)]. For obtaining leukocytes from blood, the buffy coat was obtained as previously described above, sectioned from the capillary tube and suspended in 1 × DMEM. The collected leukocytes were washed twice with 1 ×DMEM and resuspended in the same buffer for flow cytometry analyses.

### Flow cytometry of trout B cell populations and microbiota

In order to detect the IgM^+^ and IgT^+^ B cells by flow cytometry, cells suspensions from blood and gut obtained as detailed above were stained with biotinylated anti-trout IgM (clone 1.14; mouse IgG1 isotype; 1 μg/ml) and anti-trout IgT (clone 41.8; mouse IgG2b isotype; 1 μg/ml) mAbs as previously described (*6, 10*). The stained cells were detected with streptavidin-BV421 (1 μg/ml; Biolegend) and goat anti-mouse IgG2b-PE pAbs (1 μg/ml; Jackson immunoresearch). The detection of sIgs-coating on gut microbiota was performed as previously described (*10, 42*), briefly, the gut bacteria were stained with mouse anti-trout IgM (2 μg/ml) and anti-trout IgT (2 μg/ml) mAbs, then detected with goat anti-mouse IgG1-AF647 and goat anti-mouse IgG2b-PE pAbs (1 μg/ml; Jackson immunoresearch). Both steps were incubated in 1% PBS-BSA at 4 °C for 1 hour, and the stained bacteria were washed three times with 1% PBS-BSA after each staining. Mouse IgG1 and IgG2b isotypes (2 μg/ml, each; Biolegend) were used as isotype controls for primary antibody staining. To discriminate gut mucus bacteria from debris and background, all bacteria were stained with SytoBC (Invitrogen, 1 to 6000 dilutions) following manufacturer’s instructions before being applied to flow cytometer. The analyses of stained leukocyte populations and bacteria were conducted with a BD LSRFortessa™ Cell Analyzer and the data was analyzed with FlowJo software (FlowJo LLC version 10.6.1)

### Concentration measurements of trout IgM and IgT from body fluids

The previously obtained plasma (1 µl) and gut mucus (30 µl) were separated on a 4-15% Mini-PROTEAN TGX gel (BioRad) under non-reducing conditions. The gels were transferred onto a Sequi-Blot PVDF membrane (BioRad) with a Trans-Blot SD semi-dry transfer cell (BioRad). The membranes were blocked with 8% skim milk (BioRad) and incubated with mouse anti-trout IgM mAb or rabbit anti-trout IgT pAb as described in (*10*). The membranes were then detected with HRP-conjugated anti-mouse IgG (for IgM detection; GE Healthcare) or anti-rabbit IgG (for IgT detection; GE Healthcare), respectively. Immunoreactive bands were visualized using HyGLO Chemiluminescent HRP Ab Detection Reagent (Denville Scientific Products). The quantification of total IgM and IgT were conducted by scanning the western blot films and analyzing the strength of the bands with ImageJ software (NIH) and extrapolating the values to a standard curve generated using known amounts of purified IgM and IgT as previously described by us (*10, 42*).

### Analysis of bacterial translocation

Detection of translocated bacteria in gut from IgM-depleted or DSS-treated fish was performed by fluorescent *in situ* hybridization (FISH) adapting from previously described (*10*). Briefly, gut cryosections (10 µm) were fixed for 10 minutes in 4% paraformaldehyde (PFA; Wako Chemicals), permeabilized for four hours in 70% ethanol and hybridized with 5’-end-Cy5-labelled EUB338 (anti-sense probe) and 5’-end-Cy5-labelled NONEUB (control sense probe complementary to EUB338) oligonucleotide probes (Eurofins Genomics). Hybridization was performed at 41°C overnight with hybridization buffer (2 × SSC/20% formamide; Fisher) containing 3 μg/mL of the labelled probes. Slides were washed on hybridization buffer without probes followed by two more washings in PBS. Nuclei were stained with DAPI solution (1 μg/mL; Invitrogen) and slices were mounted with fluoroshield (Sigma-Aldrich) until being analyzed using a Leica DM 6000 fluorescence microscope and LAS X software.

### Endotoxin detection in trout plasma

The obtained plasma from control and depleted fish (*n* = 18-32 fish) was diluted 10-folds with endotoxin-free PBS (Millipore) and heated at 70 °C for 15 minutes. Endotoxin levels were measured by the *Limulus* amebocyte lysate (LAL) assay (Roth et al., 1990) using LAL Chromogenic Endotoxin Quantification Kit (Pierce) according to the manufacturer’s instructions. All tips, tubes and microplates used for the LAL assay were nonpyrogenic.

### Histological examination and pathology score

The whole gut from control or IgM-depleted fish was sampled and fixed in 4% PFA overnight. Samples were washed in HEPES 260 mM buffer (Gibco) and transferred to 70% ethanol prior to paraffin embedding. Blocks were sectioned at 5 μm thickness, dewaxed in xylene and stained with hematoxylin and eosin as previously done by us (*10*). A total of ten sections per fish were evaluated under a light microscope by two independent researchers in a blind fashion. The pathology score was developed based on previously reported changes in response to infections in mucosal barriers (*43, 44*). The following parameters were scored individually and added to generate the final pathology score: presence of bacterial biofilms and bacterial translocation, increased goblet cell numbers, epithelial hyperplasia, epithelial sloughing, multi-lamellar fusion, bi-lamellar fusion, distal secondary lamella necrosis, shortened secondary lamellae, presence of inflammatory infiltrates and filamental edema. Each of the selected pathological conditions were assigned with a 1 or 0 in presence or absence, respectively. The total score obtained for each analyzed section was recorded and the mean for each fish calculated.

### Dextran sulfate sodium (DSS)-induced colitis model in IgM-depleted trout

In order to assess the importance of sIgM in the gut while showing a pathological condition, we generated a dextran sodium sulfate (DSS, MP Biomedicals)-induced colitis model in trout (a common model to induce colitis in mice (*28*). Trout were injected with either anti-IgM mAb or isotype control Ab, and quarantined for 6 weeks until IgM^+^ B cells and sIgM were completely depleted from fish. One hundred microliters DSS (3% in water, w/v) was administered once a day daily by oral gavage for 10 days to either control or IgM-depleted (6 weeks since depletion) fish, and the fish were kept observed for 2 more days when DSS treatment ended. Fish were fed daily once per day 1% of biomass. Mortalities of different groups of fish were recorded from the first to twelve days since the first DSS-treatment, body weight, total length and gut length were measured from the surviving fish at the end of the experiment to evaluate *K*-factor and relative gut length.

### Ig-seq and microbiome sequencing from control and IgM-depleted fish

In order to identify taxa-specific microbiota coated by sIgM and/or sIgT, Ig-seq was performed as previously described for IgT (*10*) as an adaptation from the recently reported IgA-seq procedure (*26*). Gut mucus microbiota were obtained from 2-3 grams healthy fish as described above. Thus, microbiota was pooled in four different groups, and each pool contained microbiota for four different fish. Bacteria of each pool were stained with as described above. Therefore, 1,000,000 sIgM^+^, sIgT^+^ and sIgM/IgT^++^ bacteria were sorted per pool using a BD FACSAria II flow cytometer (BD Biosciences), while an aliquot of the whole bacteria suspension from each pool was stored as pre-sorted sample. All bacteria were kept frozen at −80°C in sucrose lysis buffer (SLB) (*45*) until further analyses. For analyzing the total gut bacteria (presort sample), four bacteria pools were used, while for the sIgM, sIgT and sIgM/IgT-coated bacteria microbiome analyses, three pools were used. All results are representative from two independent experiments. To compare the changes induced in gut microbiome due to the depletion, gut tissue from 9 IgM-depleted and 9 control fish was collected at 1, 3, 6 and 13 weeks after IgM depletion and control treatment (described above). Tissues were individually preserved in SLB buffer as previously described (*45*) and kept in −80°C until further use.

Tissue was homogenized by using sterile 3-mm tungsten carbide beads (QIAGEN) in a TissueLyser II (QIAGEN) for 5 min at a frequency of 30/sec. DNA from gut bacteria was extracted following the cetyltrimethylammonium bromide buffer method as previously described (*10, 46*). Purified DNA was then resuspended in 30 μl of deoxyribonuclease- and ribonuclease-free molecular biology grade water, measuring the concentration and purity by NanoDrop ND 1000 (Thermo Fisher Scientific). For the analysis, negative controls consisting of SLB only and positive controls consisting on a known mock bacterial community of seven bacterial species were included in each sequencing run as we previously reported (Brown 2016). Bacterial community composition was determined as by next-generation sequencing of the prokaryotic 16S rRNA (*10, 46*). Briefly, total genomic DNA for each sample was normalized to 200 ng/ul in ribonuclease-free water and amplified in triplicate using Illumina adapter fused primers that target V1 to V3 variable regions of the prokaryotic 16S rRNA sequences. Gene-specific primer sequences used were as follows: 28F, 5’-GAGTTTGATCNTGGCTCAG-3’; 519R, 5’-GTNTTACNGCGGCKGCTG-3’ (where N = any nucleotide and K = T or G). The amplification was carried out with initial activation of the enzyme (5PRIME HotMaster Taq DNA Polymerase, Quantabio) at 94°C for 90 s, followed by 33 cycles of the following: 94°C for 30 s, annealing at 52°C for 30 s, and 72°C for 90 s, and a 7-min extension cycle at 72°C with a final holding temperature of 4°C. PCR amplicons were purified using the Axygen AxyPrep Mag PCR Clean-up Kit (Thermo Fisher Scientific) as per the manufacturer’s instructions. Samples were then indexed by ligating index barcode to Illumina adapters onto the PCR amplicon using the Nextera XT Index Kit v2 Set A (Illumina). Indexed amplicon DNA concentrations in each sample were quantified, pooled, and adjusted to 200 ng/ul. Pooled samples were purified again using the Axygen AxyPrep Mag PCR Clean-up Kit and sequenced in an Illumina MiSeq platform using MiSeq Reagent Kit v3 (600 cycle) (Illumina) at the Translational Science Center at University of New Mexico Health Sciences Center.

### Microbiome sequence analysis

The sequences were analyzed by Quantitative Insights Into Microbial Ecology 2 (Qiime2, v2021.6) pipeline (*47*).We used Divisive Amplicon Denoising Algorithm (DADA2) to cluster demultiplexed sequence reads into amplicon sequence variants (ASVs) (*48*). ASVs were aligned to the latest version of the Silva 16S rDNA database (v138) in order to assign taxonomic classes (*49*). Each sample was rarefied to a sampling depth of 10500 reads before core diversity analyses were performed. Next, a core diversity analysis was conducted considering both time and treatment as main variables. Alpha diversity indices (Shannon diversity) and beta diversity measures (Weighted UniFrac distances) were generated using the QIIME2 plugin. The PCoA plots of beta diversity metrics were created using the Qiime2R package in RStudio version1.3.959 (*50*). PICRUSt2 analysis was performed in RStudio to predict the functional composition of sampled microbial communities (*51*).

### Real-time PCR for host gene expression analysis

Extraction of RNA from gut, gene expression analyses and the primers used in this study were described previously by us (*10, 41*). Real-time PCR analysis were conducted with a PowerUp SYBR green master mix (Thermo Fisher) using a 7500 Fast Real-Time PCR system (Thermo Fisher Scientific).

### Quantification of bacterial loads by qPCR

We calculated the bacterial load in each sample using 16s rRNA gene quantitative PCR as described (*52*). The prokaryotic 16S rRNA sequences were amplified in triplicate using Illumina adapter fused primers to target the V1 to V3 variable regions by gene-specific primer sequences as previously described. The qPCR reaction was prepared in a 96-well plate with 2 µl of each normalized DNA sample at 10 ng/ul, 2 ul of forward and reverse primers mix as described, 6 µl of nuclease-free water, and 10 µl of SsoAdvanced Universal SYBR Green supermix (Bio-Rad). The amplification was carried out with initial activation of the enzyme at 94°C for 90 s, followed by 33 cycles of the following: 94°C for 30 s, annealing at 52°C for 30 s, and 72°C for 90 s, and a 7-min extension cycle at 72°C with a final holding temperature of 4°C. The real-time PCR reactions were performed using the Bio-Rad CFX96 C1000 Touch detection system (Bio-Rad). To quantify 16s rRNA gene copies, each sample was aligned to a serial dilution of *E. coli* standard series of 16s rRNA gene with 10^9 to 10 V1 to V3 copies.

### Statistical analysis

Preliminary studies were conducted to the end of obtaining a representative sampling size, using power analyses to ensure a statistical power among 80 to 100%. All the sample sizes (n) and number of independent experiments are indicated in the figure captions. Fish were always sampled in a randomized way, and all experiments were at least duplicated through the paper. Fish were sampled once and were euthanized after every experiment. The histological examination of the hematoxylin and eosin micrographs to determine pathology scores in gut were performed blindly by two independent trained researchers. No data were excluded. Unpaired Student’s *t* test, One-Way ANOVA (post-hock Tuckey’s test) and Logrank test analysis were performed in GraphPad Prism (version 9.4). Significant differences were considered when p value ≤ 0.05.

**Fig. S1.**
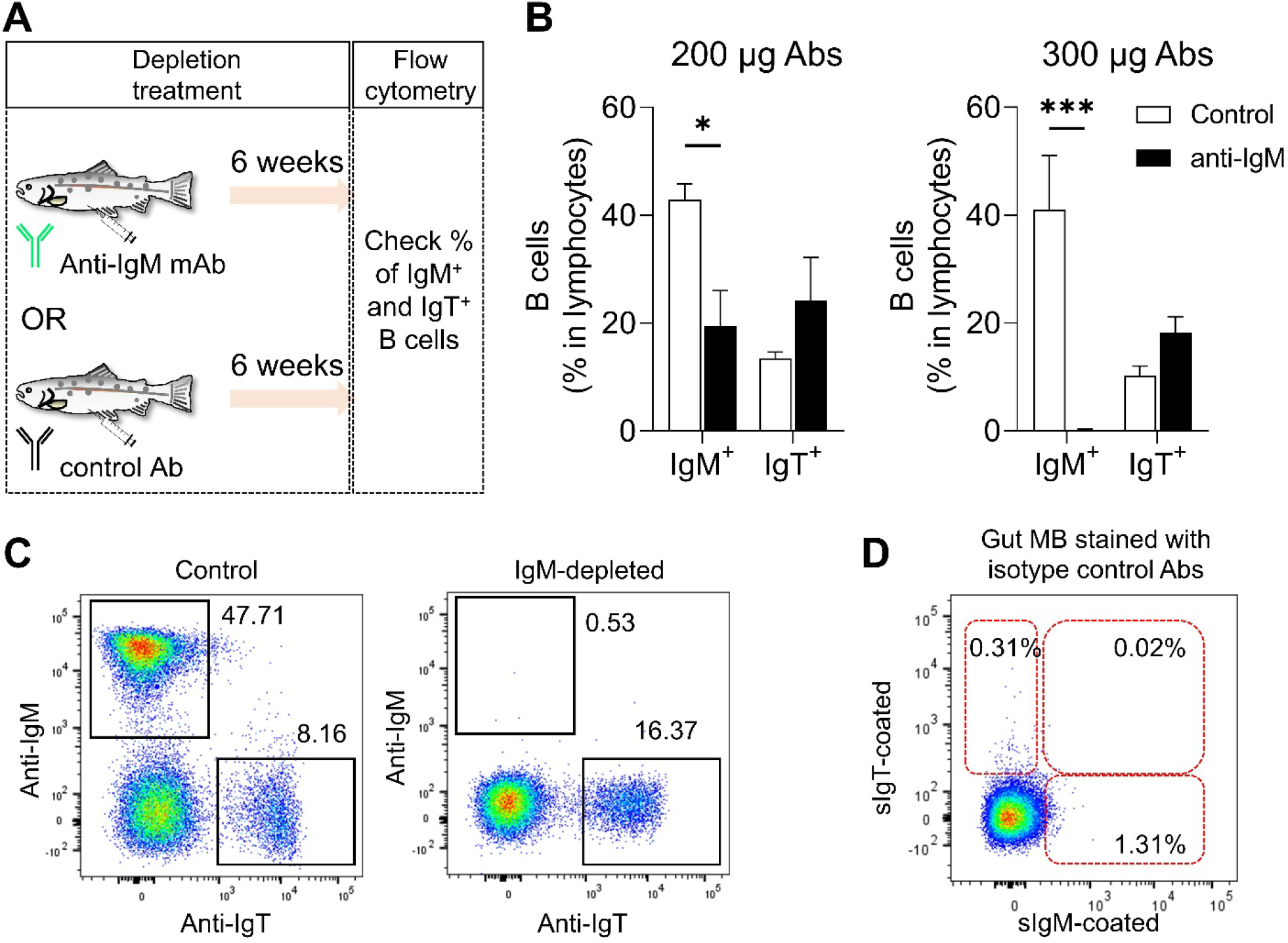
Development of the rainbow trout IgM^+^ B cell depletion model. **(A**) Scheme of the strategy used to deplete IgM^+^ B cells from trout. Fish were intraperitoneally injected with either mouse anti-trout IgM mAb or isotype control Ab. The percentages of IgM^+^- and IgT^+^ B cells from blood leukocytes of all fish was analyzed by flow cytometry at 6 weeks after IP injection. (**B** and **C**) Percentage of IgM^+^-(left bars) or IgT^+^ B cells (right bars) in blood lymphocytes from trout injected with isotype control Ab or with anti-trout IgM mAb at a dose of 200 μg (left panel) or 300 μg (right panel) per fish (*n* = 4 fish per group, means ± SEM). Statistical analysis was performed by unpaired Student’s *t*-test. \**P* < 0.05, \*\*\**P* < 0.001. (**C**) Representative dot plots of IgM^+^ and IgT^+^ B cells in blood leukocytes from control (left) and IgM^+^ B cell depleted (right) fish at 6 weeks post-treatment. Numbers adjacent to outlined areas indicate percentages of IgM^+^ B cells (top left) or IgT^+^ B cells (bottom right) in the lymphocyte population. (**D**) Dot plot representing isotype control staining of the bacteria in gut mucus from naïve fish detected by flow cytometry. Bacteria were stained with mouse IgG1 (2 μg/ml, isotype control Ab for anti-trout IgM) and IgG2b (2 μg/ml, isotype control Ab for anti-trout IgT). Numbers adjacent to outlined areas indicate percentages of microbiota stained by isotype control Abs.

**Fig. S2.**
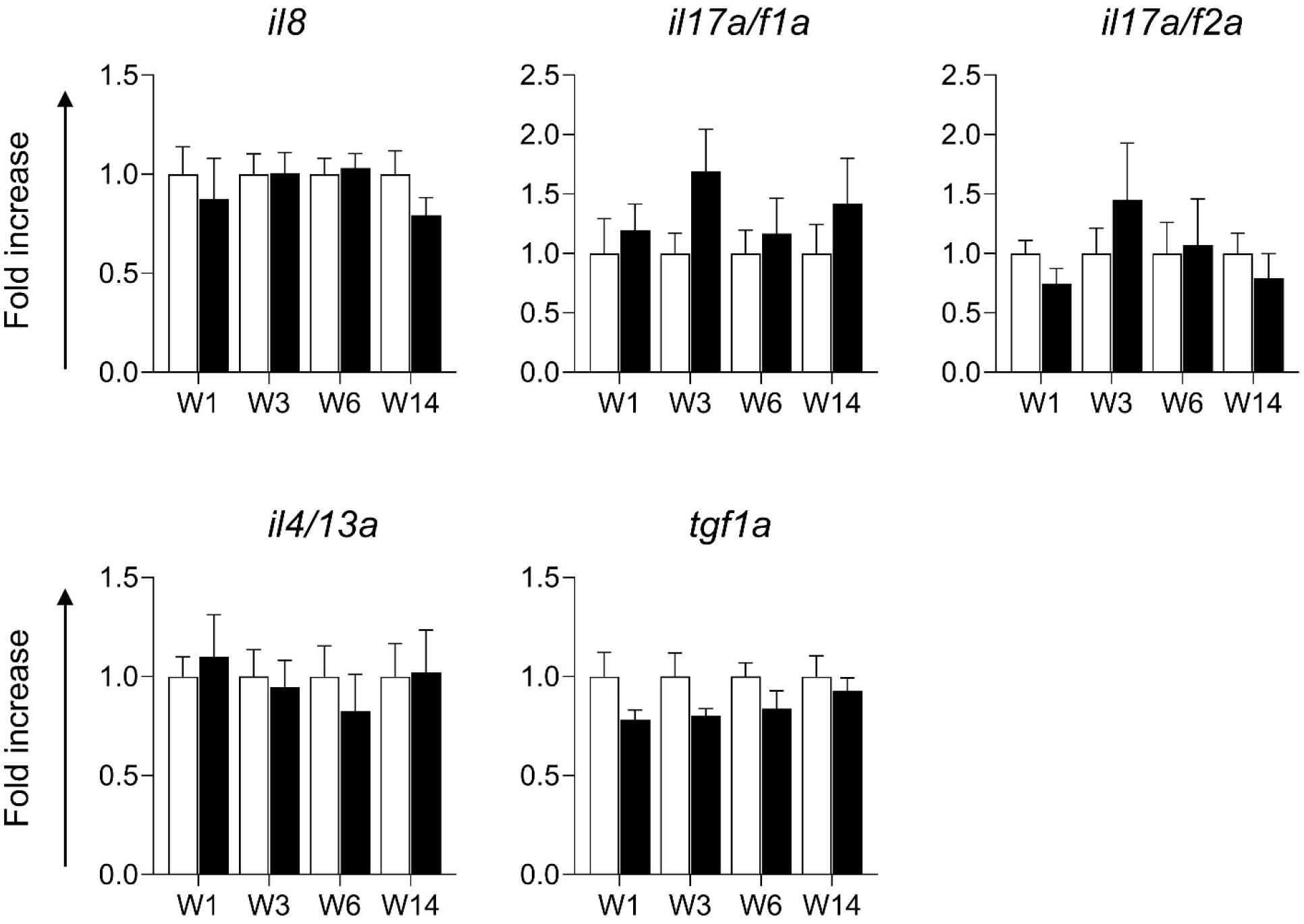
Cytokines and AMPs with unchanged expression of gene transcripts in gut tissue from control and IgM-depleted fish. Real-time PCR analysis of cytokines and antimicrobial peptides in gut tissue from control and IgM-depleted fish. Expression levels in IgM-depleted fish were normalized to those in control fish, which was set as 1 (means ± SEM; *n* = 10 fish per group). Data are representative of at least two independent experiments. Statistical analysis was performed by unpaired Student’s *t*-test.

**Table S1.** Alpha diversity metrics of the pre-sort, sIgM^+^-, sIgT^+^-, sIgM/sIgT^++^-coated and uncoated-gut bacteria from control trout. Different letters indicate significant differences (*P* <0.05).

| **Diversity Index** | **uncoated** | **Pre-sort** | **sIgM<sup>+</sup>** | **sIgT<sup>+</sup>** | **sIgM/sIgT<sup>++</sup>** |
| --- | --- | --- | --- | --- | --- |
| **Chao1** | 52.21 ± 26.44<sup>a</sup> | 51.01 ± 17.34<sup>a</sup> | 54.78 ± 22.21<sup>a</sup> | 70.57 ± 16.92<sup>a</sup> | 56.70 ± 13.28<sup>a</sup> |
| **Faith's** | 5.61 ± 1.91<sup>a</sup> | 3.76 ± 2.38<sup>b</sup> | 7.12 ± 2.89<sup>a</sup> | 6.00 ± 1.44<sup>a</sup> | 4.71 ± 1.10<sup>ab</sup> |
| **Pielou's evenness** | 0.41 ± 0.11<sup>a</sup> | 0.37 ± 0.10<sup>a</sup> | 0.46 ± 0.14<sup>a</sup> | 0.32 ± 0.08<sup>a</sup> | 3.31 ± 0.82<sup>a</sup> |
| **Shannon** | 3.08 ± 0.92<sup>a</sup> | 3.12 ± 0.88<sup>a</sup> | 2.86 ± 0.84<sup>a</sup> | 3.30 ± 0.84<sup>a</sup> | 3.30 ± 0.85<sup>a</sup> |

**Table S2.**
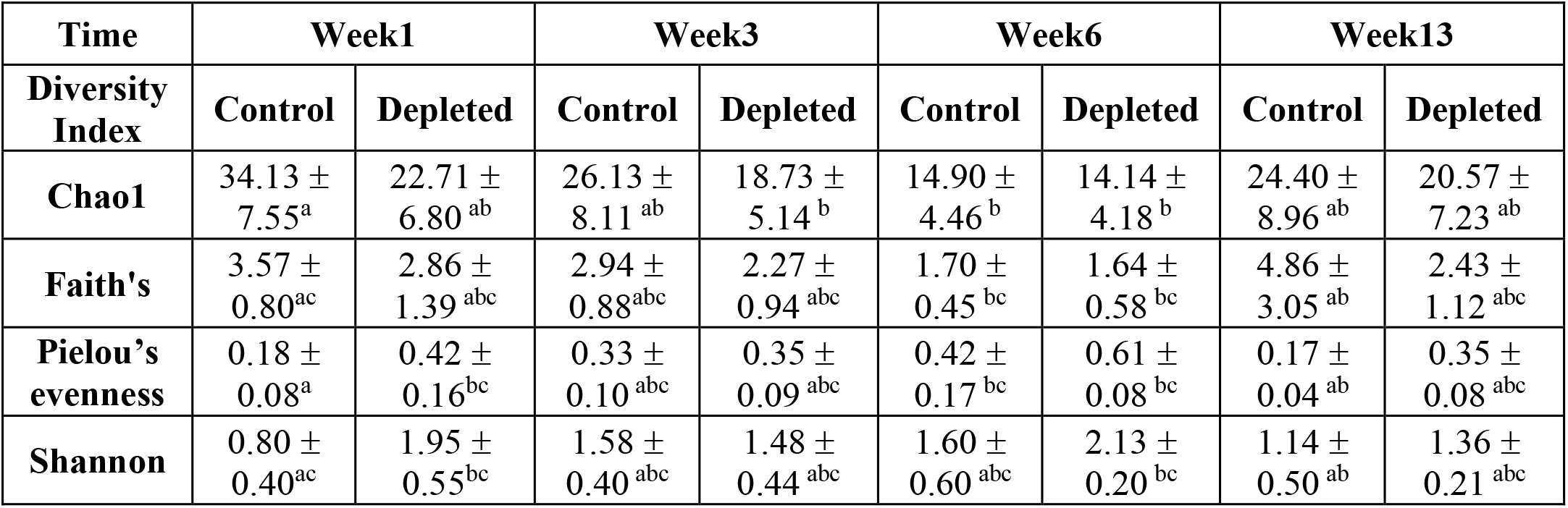
Alpha diversity metrics of the control and IgM-depleted gut microbial communities at 1, 3, 6 and 13 weeks post depletion. Different letters indicate significantly different values (P <0.05).

**Table S3.**
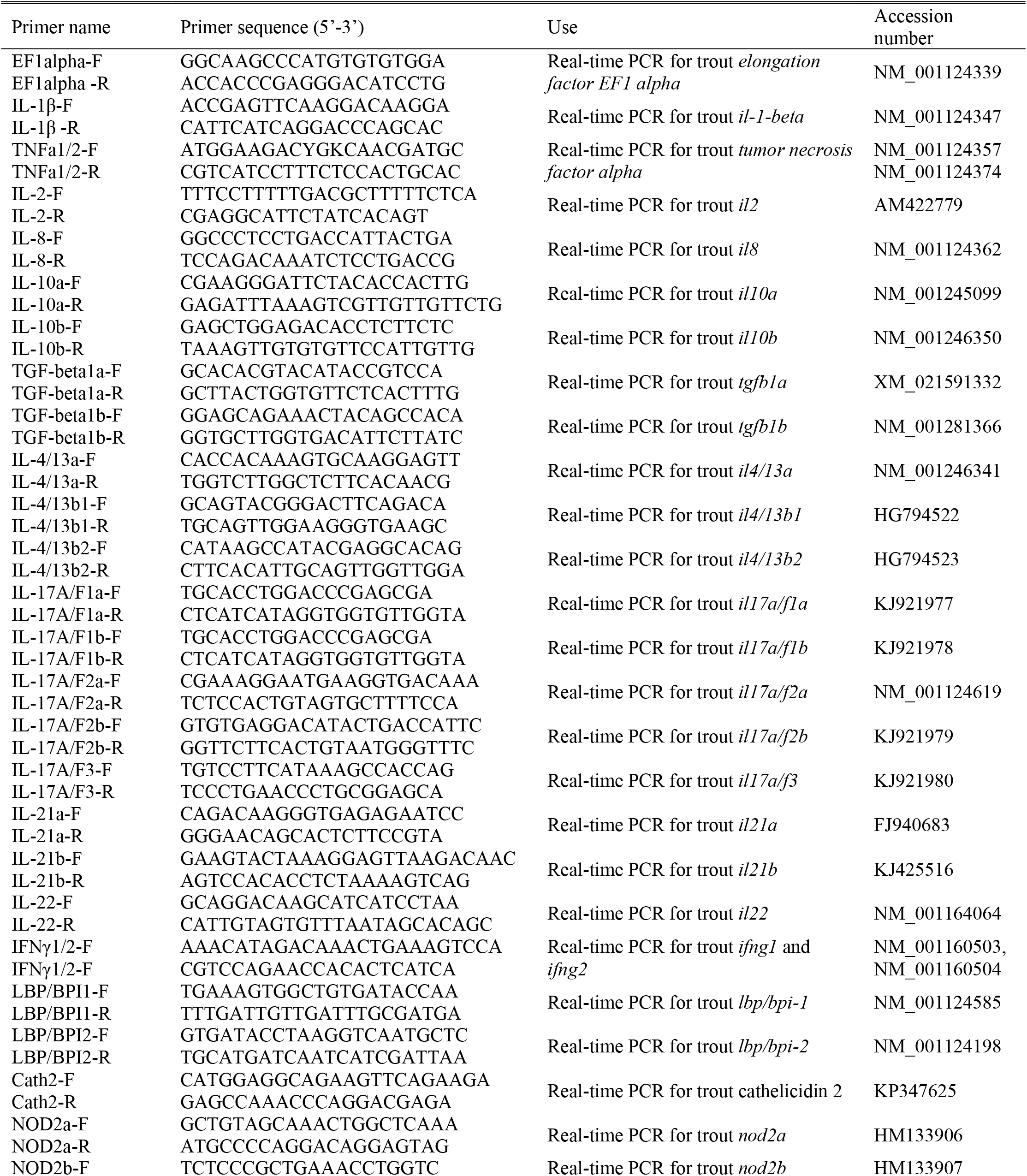
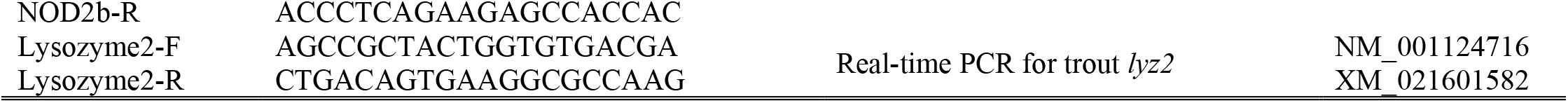
Primer sequences for real-time PCR analysis cytokines and AMPs in gut tissue from control and IgM-depleted fish at 1, 3, 6 and 13 weeks after IgM-depletion treatment.

